# Growth hormone transgenesis affects thermal tolerance in zebrafish (*Danio rerio*)

**DOI:** 10.1101/2021.05.26.445844

**Authors:** Daniela Volcan Almeida, Marcio Azevedo Figueiredo, Luis Fernando Marins

## Abstract

In fish, growth hormone (GH)-transgenesis may modify physiological mechanisms of adaptation when challenged by biotic and abiotic stressors. Thus, we evaluated whether GH overexpression can alter the thermal tolerance of adult and juvenile GH-transgenic zebrafish (*Danio rerio*). This study compared the thermal tolerance in non-transgenic (NT) and GH-transgenic (T) zebrafish exposed to 13 °C, 39 °C, or 28 °C (control) for 96 h. Mortality rate was checked every 12 h in juvenile (8 week-old) and adult males (6 month-old). Exposure to different temperatures revealed that GH overexpression increases the tolerance of transgenic juveniles exposed to 13 °C and diminishes the tolerance of juveniles and adults, when exposed to 39 °C. Additionally, we have analyzed transcriptional expression from the heat shock proteins (HSPs), which are mainly involved in the thermal tolerance mechanism. The mRNA level analysis results revealed that, under controlled conditions (28 °C), GH-transgenesis upregulates the expression of *hsp47*, *hsp70*, *hsp90a* and heat shock transcription factor (*hsf1a*) in transgenic juveniles, although the same result was not observed in transgenic adults. Exposure to low temperature did not alter the expression of any analyzed gene, both in adults and in juveniles. Exposure to 39 °C decreased the expression of all genes analyzed, in GH-transgenic adults. Furthermore, the HSP expression pattern was analyzed via hierarchical clustering. This analysis revealed two major clusters illustrating the dependency of gene changes related to age. These results indicate that the GH overexpression can alter thermal tolerance of fish, depending of age and temperature.

**Highlights:**

- GH-transgenesis increased the survival rate of juveniles at low temperature;
- High temperature is more lethal for juvenile ande adult GH-transgenic zebrafish;
- GH-transgenesis increased expression of *hsf1a*, *hsp47*, and *hsp70* genes in juvenile zebrafish;
- *hsf1a*, *hsp47*, *hsp70, hsp90a, and hsp90b* genes expression is diminished in adult zebrafish GH-transgenesis exposure at high temperature.

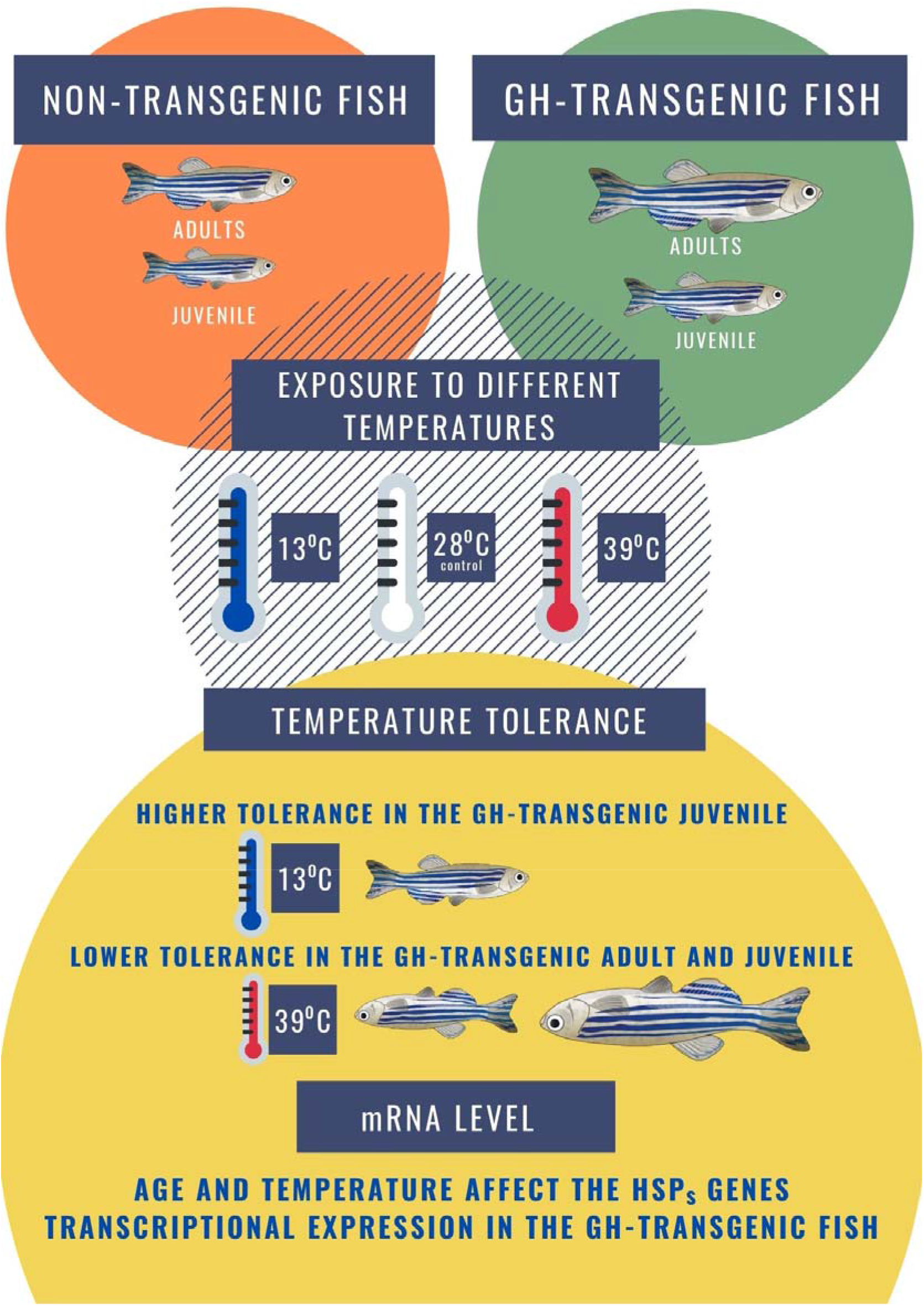

## Introduction

Growth hormone (GH) is a single-chain polypeptide that is synthesized, stored, and secreted by the adenohypophysis, which intensifies all major aspects of amino acid uptake and protein synthesis. GH-induced signaling is mediated by a group of tyrosine kinases, Janus kinases (JAKs) and their substrates, and signal transducers and activators of transcription (STATs), with the insulin-like growth factor 1 (IGF-1) being its main biological effector (Kiu and Nicholson, 2012).

GH is the principal stimulator of somatic growth, and the greatest growth is one of the most desired traits of aquaculture. For this reason, fish have been genetic manipulated, showing significantly enhanced somatic growth via muscle hypertrophy (Kuradomi et al., 2011), or hyperplasia (Hill et al., 2000).

GH-transgenesis has proved to increase the growth performance in several aquaculture fish species with impressive results, including tilapia *Oreochromis niloticus* (Rahman et al., 1998), carp *Cyprinus carpio L*. (Guan et al., 2008) and various salmonids such as coho salmon *Oncorhynchus kisutch* (Devlin et al., 2004). However, it is still necessary to assess the potential ecological consequence of growth-enhanced animals, considering that GH-transgenesis may modify physiological mechanisms of adaptation when challenged by biotic and abiotic factors. Among the different effects caused by transgenesis, it is important, both ecologically and for culture, to assess the change in tolerance to temperature. The effect of exposure to temperature was investigated in GH-transgenic coho salmon (*O. kisutch*), respectively (Chen et al. 2015). In this study, Chen and collaborators concluded that genetic modification of growth-related pathways can affect thermal performance, especially at physiological optimum or sub-critical temperatures.

To cope with the potential effects of environmental stress, fish respond with different strategies. The cellular stress response, characterized by an increase in the synthesis of heat shock protein (HSPs), is generally exhibited by all organisms (Schlesinger, 1990; Feder and Hofmann, 1999). In order to evaluate early ontogenic effects of GH transgenesis on the respiratory and cellular physiology of Atlantic salmon, Polymeropoulos et al. (2014) investigated the effects of GH transgenesis and polyploidy on metabolic, heart and ventilation rates and HSP levels after exposure to acute hypoxia in post-hatch alevins.

The HSP family comprises proteins that were originally identified based on their excess synthesis in cells exposed to elevated temperatures and were subsequently induced by exposure of cells to various stresses (Iwama et al., 1998; Basu et al., 2002). These proteins are named according to their molecular weight, emphasizing those with low molecular weight (16–47 KDa), HSP70 (68–73 KDa) and HSP90 (85–90 KDa).

Transcription of HSPs depends on the activation of a specific transcription factor, the heat shock factor (HSF-1), which binds to the heat shock element (HSE) in the HSP gene promoters (Morimoto, 1998). Activation of the HSP promoters is mediated via nuclear factors NF-IL6 and STATs, suggesting that HSP transcription can also to mediated by the JAK/STAT pathway (Stephanou et al., 1999; Madamanchi et al., 2001; Schoof et al., 2009). More recently, Jego et al. (2019) reviewed how HSPs expression is modulated by STAT signaling pathways. Nevertheless, it is necessary to understand whether JAK/STAT pathway GH-induced can be a direct modulator of HSP expression, since the excessive GH presumably affects the (HSPs) expression (Deane and Woo, 2005; Deane and Woo, 2010).

Considering all the previously mentioned aspects, our study aimed to test the hypothesis that GH overexpression can alter the thermal tolerance of adult and juvenile transgenic zebrafish. As a result, we exposed juveniles and adults from a GH-transgenic zebrafish lineage (produced as described in Figueiredo 2007) to thermal stress. Tolerance to low and high temperature was related to *hsp* transcriptional expression.

## Material and Methods

### Fish

Transgenic and non-transgenic zebrafish were obtained from crosses between non-transgenic females and hemizygous GH-transgenic males from lineage F0104. The lineage F0104, used in this study, were produced by Figueiredo et al. 2007. Freshly fertilized eggs were collected for microinjection and transgenic zebrafish produced using two transgenes containing the carp (*Cyprinus carpio*) β-actin promoter driving the expression of either the *A. Victoria* GFP gene (pcβA/GFP plasmid) or the marine silverside fish (*Odonthestes argentinensis*) growth hormone (msGH) cDNA (pcβA/msGH plasmid).

The fish underwent fasting during temperature treatment period (96 h for thermal tolerance assay; 12 h and 2 h for expression gene assay, in 13 °C and 39 °C, respectively). All experimental procedures were performed as indicated by the Ethics for Animal Use Committee from the Federal University of Rio Grande, where this work was carried out (23116.005574/2013-03).

### Temperature tolerance of GH-transgenic zebrafish

Juvenile (8 week-old) and adult males (6 month-old) of non-transgenic and transgenic zebrafish were exposed to 13 °C, 39 °C, or 28 °C (control) for 96 h in an incubator. To avoid rapid changes of water temperature, fish were first put into each incubator at a temperature of 28 °C and temperature was increased or decreased in a continuous linear rate of ±3 °C·h^−1^. Mortality rate was checked every 12 h. Each experimental temperature was tested in triplicate containing 10 fish in each experimental unit (one fish per liter for adults and five fish per liter for juveniles). A total of 60 adult fishes (30 non-transgenic and 30 transgenic) and 60 juvenile fishes (30 non-transgenic and 30 transgenic). The experimental tank were continuously aerated and parameters such as temperature, pH, nitrate concentration, and photoperiod (14h light/ 10h dark) were daily monitored. Death criteria included the lack of both operculum movement and twitching reflex upon prodding with a needle.

### mRNA level

All experimental groups were exposed to 13 °C and 39 °C for 12 and 2 h, respectively. The exposure times were those considered as sublethal according to data from the thermal tolerance experiment. Control animals also exposed to 28 °C, and sampled at the same experimental condition as the treated groups (12 h for control of 13 °C and 2h for control of 39 °C). mRNA level was determined in livers from zebrafish of all groups (*n* = 6 per group). The liver, the hub of metabolism, was selected as the target tissue. Livers were pooled in juveniles (3 independent pools per treatment, 18 fish in total per group). For adults were collected liver individually (*n* = 6 fish per group). Total RNA was prepared with Trizol reagent (Invitrogen, Brazil) and purified with RNAse free DNAse I (Invitrogen, Brazil), following the manufacturer’s protocol. For cDNA synthesis, 2 μg of total RNA was reverse-transcribed using a High Capacity cDNA Reverse Transcription Kit (Invitrogen, Brazil). The obtained cDNA was used as a template for gene amplification using specific primers. Gene-specific primers (Table 1) were designed based on sequences available in GenBank using the Primer Express 3.0 software (Applied Biosystems, Brazil). Previously, the PCR amplification efficiency of each primers pair was evaluated by serial dilutions reactions where the efficiency of reactions showed appropriate parameters. Quantitative

**Table 1.**
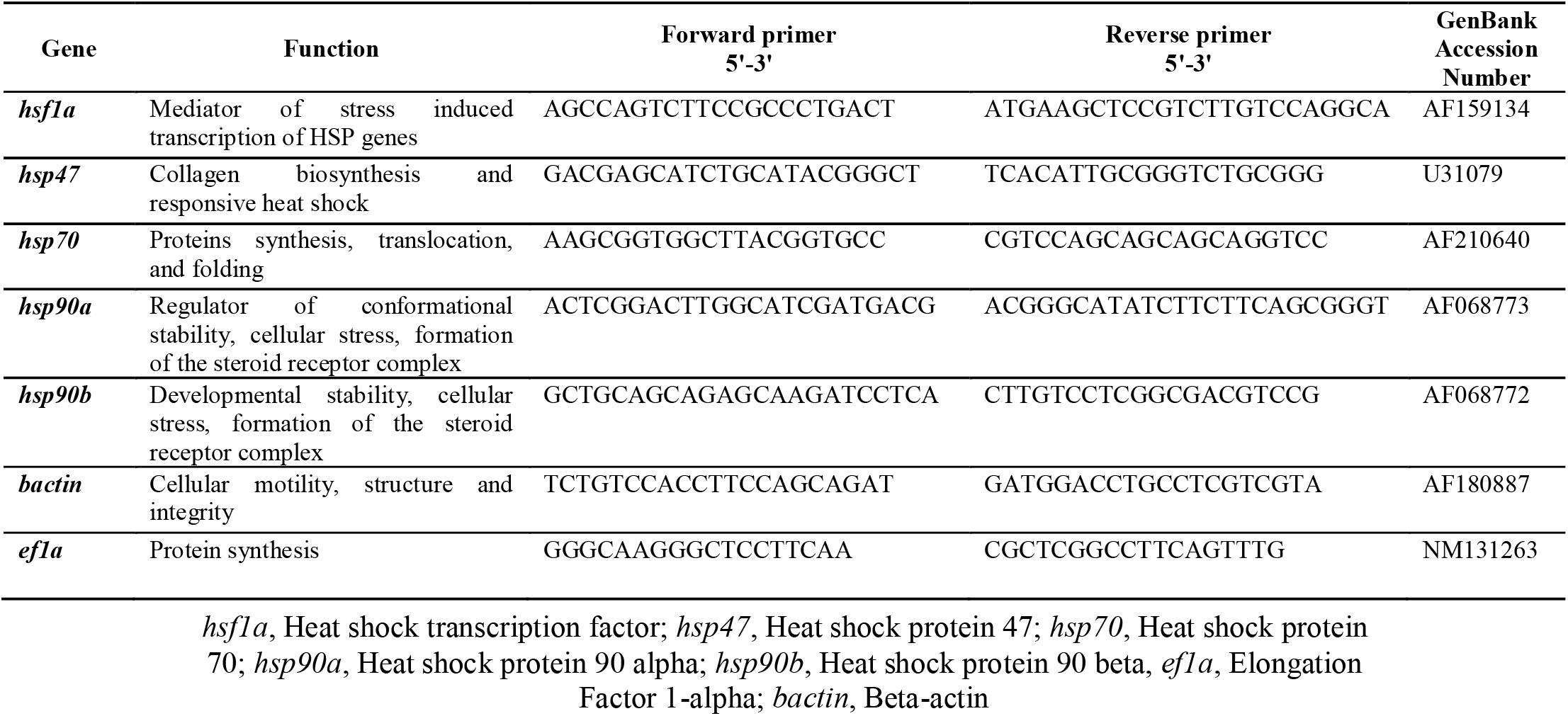
Primer sequences used for RT-qPCR analysis of mRNA expression in zebrafish (*Danio rerio*)

PCR was performed with an ABI 7500 System (Applied Biosystems, Brazil) using Platinum SYBR Green qPCR SuperMix-UDG (Invitrogen, Brazil). The PCR program consisted of 40 cycles of 95 °C for 15 s and 60 °C for 30 s after an initial cycle of 50 °C for 2 min and 95 °C for 2 min. The synthesized cDNA (1 μL) was used for the qPCR reactions in a total volume of 10 μL and a final concentration of 200 nM per each primer. Triplicates were run for each sample. Melting curve analysis was performed during qPCR to identify the presence of primer dimers and analyze the specificity of the reaction Expression levels of target genes were normalized using the expression of two housekeeping genes, elongation factor 1a (*ef1a*) and β-actin (*bac*). These genes were choosen as reference genes after presented stability when tested with geNorm applet (Vandesompele et al. 2002).

### Statistical analyses

Fish survival curve (%) among treatments was performed by the log-rank test, where significant differences were considered at p<0.05. The data were analyzed using GraphPad Prism® following the instructions of GraphPad Statistic Guide for comparisons of survival curves. The accumulative survival at 12 h was analyzed by Student’s t test, comparing non-transgenic and transgenic, in each temperature separately. Relative mRNA level were analyzed by Relative Standard Curve Method between the transgenic and non-transgenic groups, in each temperature. Expression levels of target genes were normalized by normalization factor using the expression of two housekeeping genes, elongation factor 1a (*ef1a*) and β-actin (*bac*). The values obtained were statistically analyzed by Student’s t test. Hierarchical clustering of gene expression and heat maps were produced using PermutMatrix, using McQuitty’s method (Caraux and Pinloche, 2005).

## Results and discussion

To evaluated whether GH overexpression can alter the thermal tolerance of adult and juvenile GH-transgenic zebrafish (*Danio rerio*), our compared the thermal tolerance in non-transgenic (NT) and GH-transgenic (T) zebrafish exposed to 13 °C, 39 °C, or 28 °C (control) for 96 h. Mortality rate was checked every 12 h in juvenile (8 week-old) and adult males (6 month-old). According the log-rank test, in juveniles, exposed at 13 °C, a significant difference of survival curve was observed, indicating that transgenic juveniles are more tolerant to low temperature than non-transgenic. The higher tolerance of GH-transgenic fish at 13°C may be related to the metabolism of these animals. The metabolic rate is diminished in lower temperature, and changes in oxygen and energy delivery as kinetic processes respond directly to temperature. Low temperature reduces resting heart rate and cardiac output in fish and consequently decreases metabolism (Gamperl, 2011). However, the overexpression of GH increases the metabolic rate in GH transgenic zebrafish (Rosa et al., 2008), this increase may alleviate adverse effects.

For adults group, no significant effect was observed on the survival curve under low temperature (13 °C). Similarly, no significant difference was observed in high temperature (39 °C) between the survival curve of non-transgenic and transgenic adults and juveniles (Fig. 1). However, if we analyze only the mortality in 12 h of exposure to 39 °C, there are a significant effect (*t* test, p<0.05) on the final survival of transgenic juveniles and adults (Fig. 1). This finding, at 12 h, indicating thermal tolerance difference between the transgenic and non-transgenic juveniles and adults exposed at 39 °C (*t* test realized in point 12 h, p<0.05). Thus, the exposure to different temperatures has indicated that GH overexpression can alter the thermal stress tolerance depending on age and temperature in GH-transgenic zebrafish.

**Figure 1.**
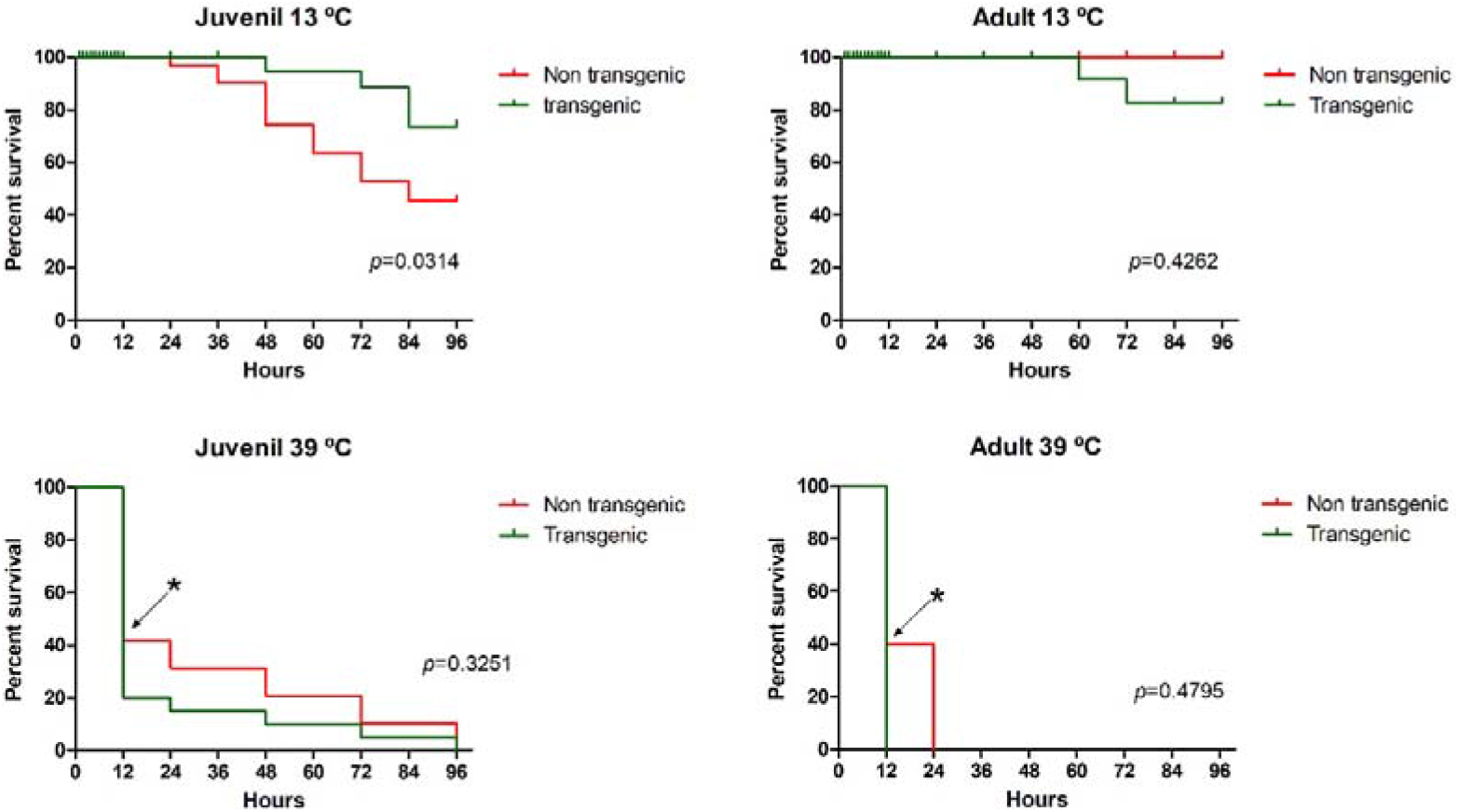
Survival percentage of juvenile and adult zebrafish (*Danio rerio*) exposed to 13 °C and 39 °C for 96 h. Data are presented as means ± SEM (*n* = 10 for each replicate; *n* = 3 replicates). *p* values represent curve comparison by Log-rank (Mantel-Cox) test. Asterisk (*) indicate significant different between juveniles NT and T in 12 h (*t* test, *p* < 0.05).

In contrast to the results obtained at low temperature, both juvenile and adult transgenic fish were less tolerant to high temperature, where most *hsps* were downregulated. Vergauwen et al. (2010) have reported that zebrafish exposed to high temperatures present diminished protein and lipid content in liver, which evinces that, during warm acclimation, the increased metabolic rate results in increased energy expenditure.

Unlike low temperatures, high temperatures tend to increase the metabolic rate (Gamperl, 2011) of fish. Thus, when GH-transgenic zebrafish were exposed to 39 °C, they had the adverse effect of your high metabolic rate potentiated by high temperature, causing transgenic fish to die before death non-transgenic fish.

With an aim to explain the observed alteration of thermal tolerance in transgenic groups, we analyzed the *hsps* gene expression profile. In juveniles, under control conditions (Fig. 2), GH-transgenesis promoted a significant rise of 10.6-, 14.3-, 9.5-, and 15.8-fold in the expression of *hsf1a*, *hsp47*, *hsp70*, and *hsp90a*, respectively. In this condition, similar effect was not observed in adults, wherein only the expression of *hsp90a* was altered (decrease by 72%). At low temperature (13 °C; Fig. 2), *hsp47* expression increased in the juveniles (3.9-fold induction), whereas *hsp90b* genes were downregulated (decrease of 60%). In adults, only *hsp90a* gene was upregulated by 1.7-fold in the transgenic fish.

**Figure 2.**
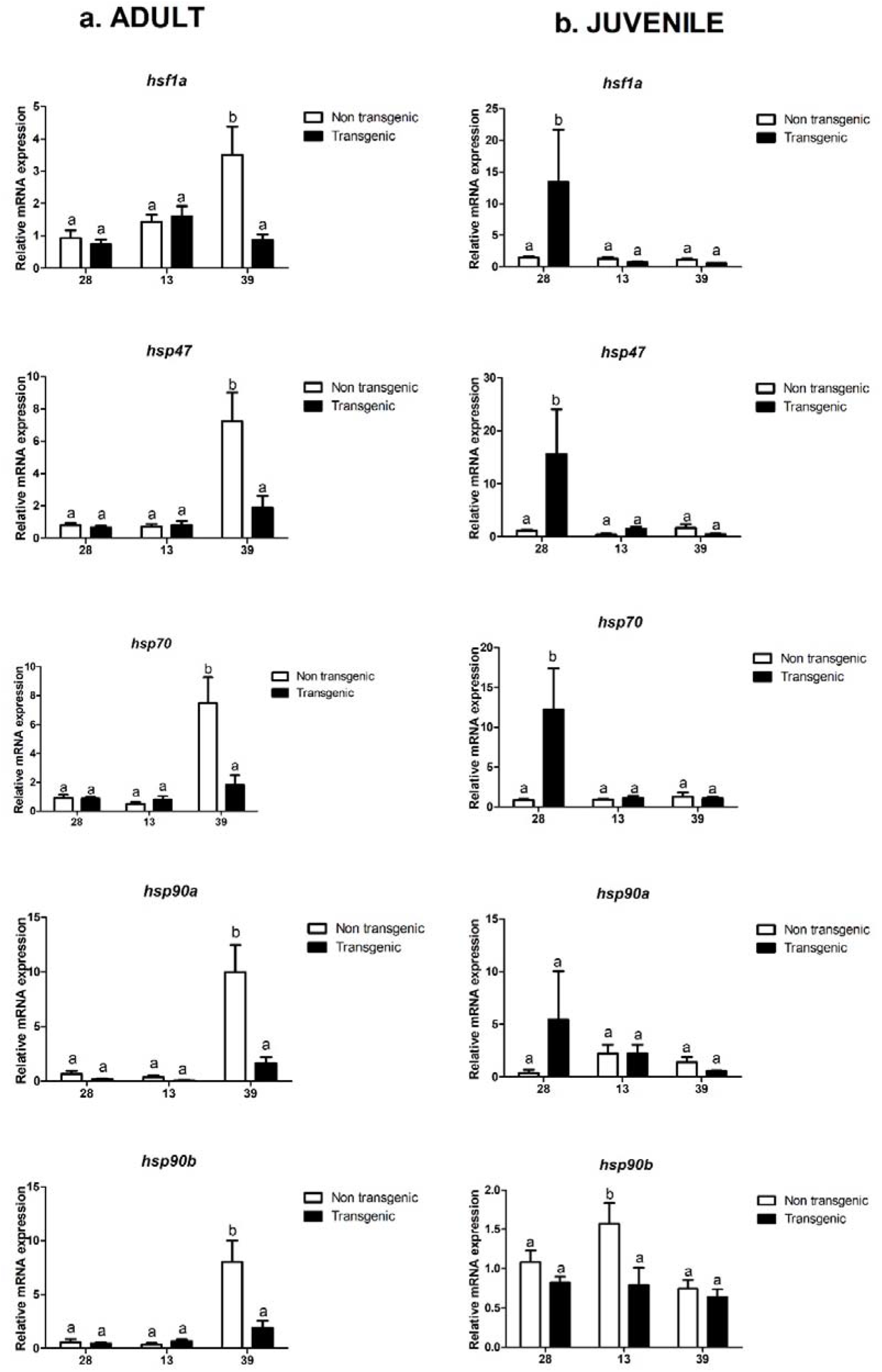
Relative expression of HSP family genes in livers of non-transgenic (white bars) and GH-transgenic (checkered bars) zebrafish (*Danio rerio*) kept under control condition (28 °C), low temperature (13 °C) and high temperature (39 °C). mRNA level was normalized by the expression of the elongation factor 1 alpha (*ef1a*) and β-actin (*bac*) gene. Data are presented as median ± SEM (*n* = 6 per group). Livers of juveniles were pooled (3 livers per *n*) and liver of adults were collected individually. Significant differences (*p* < 0.05, *t* test) in mRNA levels are denoted by an asterisk (*). *hsf1a*, Heat shock transcription factor; *hsp47*, Heat shock protein 47; *hsp70*, Heat shock protein 70; *hsp90a*, Heat shock protein 90 alpha; *hsp90b*, Heat shock protein 90 beta.

Exposure to high temperature (Fig. 2) decreased the expression of a majority of genes in both transgenic groups. *hsf1*a, *hsp47*, and *hsp90a* were downregulated in the transgenic juveniles (49, 74, and 64 %, respectively). Whereas *hsf1a*, *hsp47*, *hsp70*, *hsp90a*, and *hsp90b* were downregulated in the transgenic adults (49, 58, 53, 71, 47 %, respectively).

Under normal rearing conditions (28 °C), transgenic juveniles presented higher expression in most HSPs compared to the non-transgenic ones. The ability of organisms to face a stressful situation is associated with its physiological condition, in particular, with its energy status (Tseng and Hwang, 2008). In transgenic juveniles, their entire energy might have been deviated to growth, stimulated by the excess of GH, resulting in increased protein synthesis. The folding of already synthesized proteins depends on the action of molecular chaperones. Thus, the highest growth rate found in transgenic zebrafish (Figueiredo et al. 2007, Studzinski et al. 2009) could trigger the HSP transcription. In contrast, in transgenic adult animals, no increase in HSP transcription (in 28 °C) was observed by transgenesis effect. One possible explanation is that, unlike most fish, the zebrafish exhibit determinate muscle growth, with growth plateau after the juvenile phase (Biga and Goetz, 2006), decreasing the need for increased HSP synthesis.

In addition to the important role of HSP chaperones in multiple cellular processes such protein translocation and protein folding, they are involved in regulation of the stress response (Lindquist, 1986; Lindquist and Craig, 1988). In fish, HSPs are clearly proven to be involved in thermal responses of fish (Iwama et al., 1998). Considering that overexpression of GH led to increased expression of HSP in juvenile, our hypothesis is that GH overexpression can alter the thermal tolerance of juvenile GH-transgenic zebrafish. This hypothesis was only confirmed for the lowest temperature and, at the transcriptional expression level, only *hsp47* was induced. HSP47 plays a key role in collagen biosynthesis and its structural assembly (Mala and Rose, 2010), and its relation with heat stress has been already demonstrated (Murtha and Keller, 2003). Nevertheless, few studies have evaluated the role of HSP47 in cold tolerance response of fish.

The hierarchical clustering of gene expression and the heat map are presented in Figure 3. The analysis simultaneously considered all genes and all tested temperatures for juveniles and adults groups, emphasizing two major clusters. Cluster I comprises a majority of *hsp* genes from juvenile fish, whereas cluster II majorly includes HSP genes from adults. This heat map demonstrates that age and temperature affect the *hsp* gene transcriptional expression. Similarly, Murtha and Keller (2003) reported that age-related changes were evident on transcriptional expression level of *hsps* in zebrafish. This result emphasizes an age/temperature-related HSP expression pattern.

**Figure 3.**
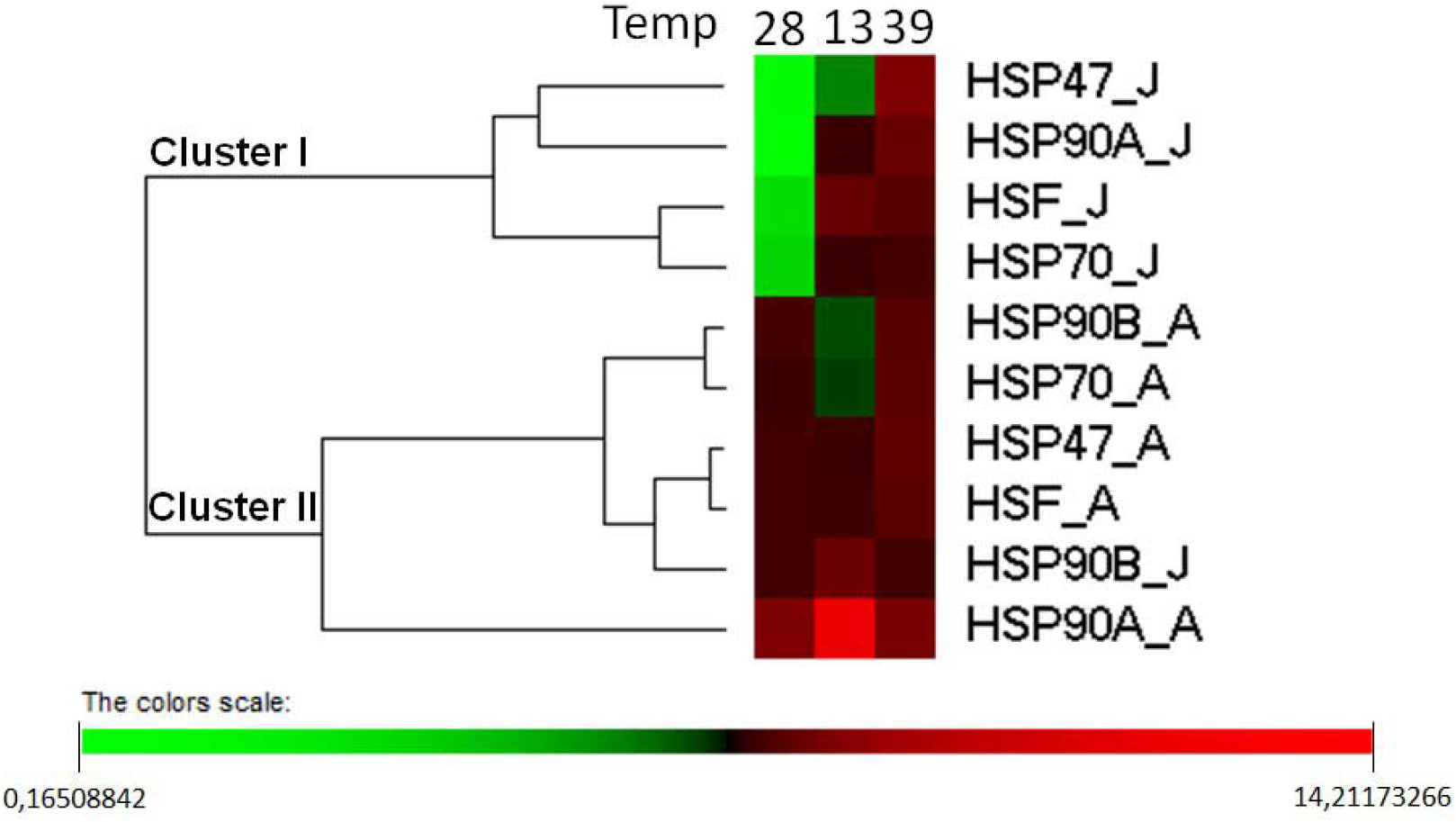
Hierarchical cluster analysis of heat shock protein (HSP) expression. The relative fold change in gene expression for genes *hsf1a*, *hsp47*, *hsp70*, *hsp90a* and *hsp90b* were compared in GH-transgenic zebrafish normalized by non-transgenic zebrafish, exposed to different temperatures (28 °C; 13 °C and 39 °C). Red indicates the downregulation of gene expression, and green refers to the upregulation of gene expression. *hsf1a*, Heat shock transcription factor; *hsp47*, Heat shock protein 47; *hsp70*, Heat shock protein 70; *hsp90a*, Heat shock protein 90 alpha; *hsp90b*, Heat shock protein 90 beta.

In summary, we assume that the effect of GH excess and thermal stress is crucial in tolerance to different temperatures in zebrafish. The results of the present study emphasize the importance of gathering knowledge concerning the collateral effects of GH-transgenesis for different environmental stressors.

## Acknowledgments

This work was supported by Brazilian CNPq (Conselho Nacional de Desenvolvimento Científico e Tecnológico) and CAPES (Coordenação de Aperfeiçoamento de Pessoal de Nível Superior). L.F. Marins is a research fellow from CNPq/Brazil (Proc. number 305928/2015-5).

## Notes

### Competing Interest Statement

The authors have declared no competing interest.

## References

Basu N, Todgham AE, Ackerman PA, Bibeau MR, Nakano K, Schulte PM, Iwama GK. 2002. Heat shock protein genes and their functional significance in fish. Gene 295: 173–183.

Biga PR, Goetz FW. 2006. Zebrafish and giant danio as models for muscle growth: determinate vs. indeterminate growth as determined by morphometric analysis. American Journal of Physiology-Regulatory, Integrative and Comparative Physiology. 291(5): 1327–37.

Caraux G, Pinloche S. 2005. PermutMatrix: a graphical environment to arrange gene expression profiles in optimal linear order. Bioinformatics. 21(7):1280–1.

Deane, E.E., Woo, N.Y.S. Growth hormone attenuates branchial HSP70 expression in silver sea bream. 2010. Fish Physiol Biochem 36: 135–140.

Deane EE, Woo NY. Growth hormone increases hsc70/hsp70 expression and protects against apoptosis in whole blood preparations from silver sea bream. 2005. Ann N Y Acad Sci. 1040: 288–92.

Devlin RH, Biagi CA, Yesaki TY. 2004. Growth, viability and genetic characteristics of GH transgenic coho salmon strains. Aquaculture: 1–26.

Figueiredo MDA, Lanes CFC, Almeida DV, Marins LF. 2007. Improving the production of transgenic fish germlines: in vivo evaluation of mosaicism in zebrafish (*Danio rerio*) using a green fluorescent protein (GFP) and growth hormone cDNA transgene co-injection strategy. Genetics and Molecular Biology 30: 31–36.

Gamperl AK. 2011. Integrated Responses of the Circulatory System to Temperature. In: Farrel AP (Ed.) Encyclopedia of Fish Physiology: from Genome to Environment. 1: 1197–1205

Guan B, Hu W, Zhang T, Wang Y, Zhu Z. 2008. Metabolism traits of “all-fish” growth hormone transgenic common carp (*Cyprinus carpio L*.). Aquaculture 284: 217–223.

Iwama GK, Thomas PT, Forsyth RB, Vijayan MM. 1998. Heat shock protein expression in fish. Reproductive Biology and Endocrinology 8: 35–56.

Kiu H, Nicholson SE. 2012. Biology and Significance of the JAK/STAT Signalling Pathways. Growth Factors 30(2): 88–106.

Lindquist, S. 1986. The heat-shock response. Annual Review of Biochemistry 55: 1151–1191.

Lindquist, S. and Craig, E.A. 1988. The heat-shock proteins. Annual Review of Genetics. 22: 631–677.

Madamanchi NR, Li S, Patterson C, Runge MS. 2001. Reactive Oxygen Species Regulate Heat-Shock Protein 70 via the JAK/STAT Pathway. Arteriosclerosis, Thrombosis, and Vascular Biology 21: 321–326.

Mala JGS, Rose C. 2010. Interactions of heat shock protein 47 with collagen and the stress response: an unconventional chaperone model? Life sciences 87: 579–86.

Morimoto RI. 1998. Regulation of the heat shock transcriptional response: cross talk between a family of heat shock factors, molecular chaperones, and negative regulators. Genes & Development 12: 3788–3796.

Murtha J, Keller ET. 2003. Characterization of the heat shock response in mature zebrafish (*Danio rerio*). Experimental Gerontology 38: 683–691.

Rahman MA, Rohan M, Hala A, Alan S, Maclean N. 1998. Expression of a novel piscine growth hormone gene results in growth enhancement in transgenic tilapia (*Oreochromis niloticus*). Transgenic Research 7: 357–369.

Rosa CE, Figueiredo MA, Lanes CFC, Almeida DV, Monserrat JM, Marins LF (2008) Metabolic rate and reactive oxygen species production in different genotypes of GH-transgenic zebrafish. Comparative Biochemistry Physiology B 149:209–214

Studzinski AL, Almeida DV, Lanes CF, Figueiredo MA, Marins LF (2009) SOCS1 and SOCS3 are the main negative modulators of the somatotrophic axis in liver of homozygous GH-transgenic zebrafish (*Danio rerio*). General Comparative Endocrinology 161:67–72

Schoof N, von Bonin F, Trümper L, Kube D. 2009. HSP90 is essential for Jak-STAT signaling in classical Hodgkin lymphoma cells. Cell communication and signaling: CCS 7: 17.

Stephanou A, Isenberg DA, Nakajima K, Latchman DS. 1999. Signal transducer and activator of transcription-1 and heat shock factor-1 interact and activate the transcription of the Hsp-70 and Hsp-90beta gene promoters. The Journal of biological chemistry 274: 1723–8.

Tseng YC, Hwang PP. 2008. Some insights into energy metabolism for osmoregulation in fish. Comparative biochemistry and physiology. Toxicology & pharmacology: CBP 148: 419–29.

Vandesompele J, De Preter K, Pattyn F, Poppe B, Van Roy N, De Paepe A, Speleman F. 2002. Accurate normalization of real-time quantitative RT-PCR data by geometric averaging of multiple internal control genes. Genome Biology. 3(7): 3401–11

Vergauwen L, Benoot D, Blust R, Knapen D. 2010. Long-term warm or cold acclimation elicits a specific transcriptional response and affects energy metabolism in zebrafish. Comparative biochemistry and physiology. Part A, Molecular & integrative physiology 157: 149–57.

